# Single-particle tracking of Nucleotide Excision Repair proteins inside living bacteria

**DOI:** 10.1101/293910

**Authors:** Alicja Piotrowska, Mathew Stracy, Pawel Zawadzki

## Abstract

Single-particle tracking (SPT) combined with Photoactivated Localization Microscopy (sptPALM) provides an opportunity to perform complex molecular biology experiments inside living cells. By tracking the motion of DNA repair proteins *in vivo*, information can be extracted not only about their diffusion, but also about the kinetics and spatial distribution of DNA binding^1–3^. From a methodological point of view, a Total Internal Reflection Microscope equipped with a sensitive detector (usually an EM-CCD camera^4^) is commonly used, allowing detection of individual fluorophores. The signal from individual emitters can be analysed and the position of a given fluorophore established with high accuracy (up to a single nm) by Gaussian fitting. To determine the mobility of each fluorophore, the positions of individual molecules are linked into trajectories over multiple frames using a tracking algorithm^5^.

---

Since most proteins in bacteria are present at a copy number, which is too high to resolve individual fluorophores, photoactivable fluorescent proteins (PAFPs i.e. PAmCherry) can be used, allowing the level of active fluorophores to be controlled (e.g. by varying the intensity of a 405 nm photoactivation laser) such that ~1 fluorophore is active per cell. This allows for the consecutive observation of all labelled proteins^1–3^. As an alternative to PAFPs, protein tags (i.e. HaloTag), which bind organic fluorophores provided externally can also be used. Once a functional fusion of the protein of interest to a fluorescent label has been constructed, the experiment can begin.

One example of the power of sptPALM was a study of the Nucleotide Excision Repair (NER) pathway in *Escherichia coli^6^*. Fusions of UvrA and UvrB - the proteins that initiate NER, to PAmCherry were introduced into the chromosome and their behaviour was studied in cells, before and after DNA lesions were induced by exposure to UV light. A movie was recorded with 15 ms frame rate and the positions of the fluorophores were localised in each frame and linked into trajectories. For each trajectory, an apparent diffusion coefficient was calculated based on the distance between subsequent localisations^5^ (Fig1. A). Molecules bound to DNA showed a minimal change in position, whereas freely diffusing molecules showed large displacements between consecutive frames. The different populations of UvrA and UvrB molecules were quantified by fitting the distribution of apparent diffusion coefficients from tens of thousands of molecules (Fig1. B, C).

**Figure 1.**
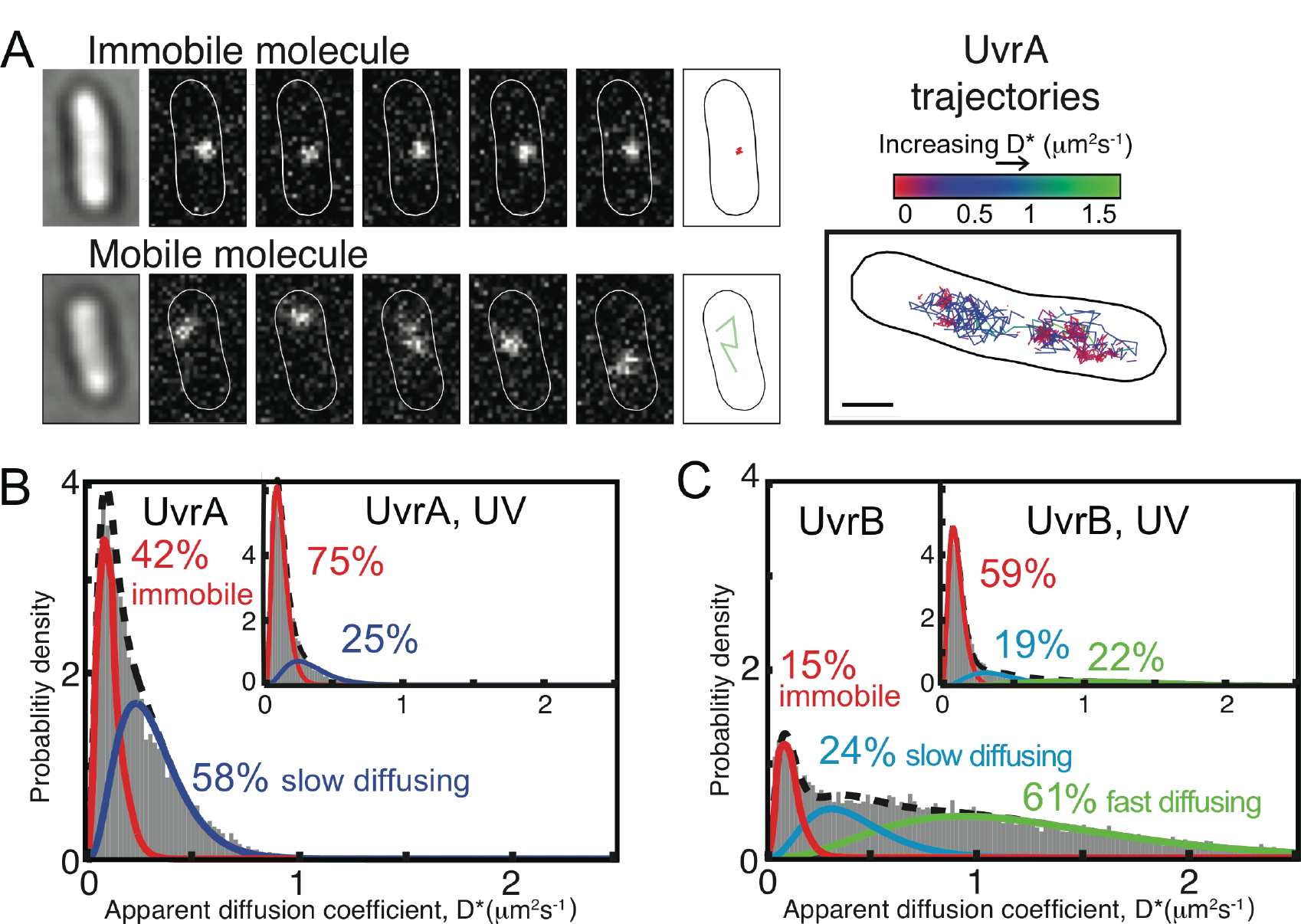
In vivo characterization of UvrA and UvrB proteins. (**A**) The example image of a single immobile UvrA-PAmCherry molecule localized and tracked at 15ms exposures within five consecutive frames (top) and the example image of five consecutive frames showing fast diffusing UvrB-PAmCherry molecule (bottom). On the right, example cell is shown with multiple trajectories recorder for many individual UvrA molecules (**B**) Distribution of apparent diffusion coefficients (D*) of tracked UvrA molecules, fitted with a two species model: first immobile, DNA-bound population (~42%) and second mobile population of slowly diffusing molecules (~58%). (Inset) The distribution of D* values of tracked UvrA molecules after exposure to 50 J m^-2^ ultraviolet light (UV). (**C**) Distribution of D* values of tracked UvrB molecules, fitted with a three species model established that ~15% of UvrB molecules were immobile, ~24% diffusing slowly and ~61% fast diffusing. (Inset) The distribution of D* values of tracked UvrB molecules after exposure to ultraviolet light (UV).

UvrA was found to bind DNA stably for ~3 s (~40% of molecules) or interact with DNA transiently (low ms range) (Fig1. B). It was proposed that the transient protein-DNA binding is a part of the initial DNA search process, whereas longer binding is a damage verification step. On the other hand, UvrB showed very different behaviour, with the majority of UvrB molecules freely diffusing throughout the cytoplasm (Fig1. C). Cell exposure to UV light caused the recruitment of most UvrA and UvrB molecules to DNA (75% and 60% of molecules, respectively) to repair UV-induced lesions. These sptPALM experiments showed that, in contrast to some historical *in vitro* experiments, UvrA and UvrB rarely form a complex in solution; instead, UvrA is a DNA damage sensor, recruiting UvrB to DNA only after damage detection. Furthermore, by using catalytic mutants of UvrA, it was possible to decipher the roles of the two ATP binding sites present in each UvrA molecule, showing that cooperative action in both sites is necessary to recruit UvrB to DNA damage sites^6^.

## Cautionary notes

Previously, one of the key factors preventing the wider adoption of SPT and sptPALM in microbiology has been limited access to the sophisticated equipment and custom-written data analysis tools required for imaging single molecules. However, as these techniques increase in popularity, commercial single-molecule microscopes are becoming more affordable and data analysis tools for SPT are becoming more available^7^, opening access to this technique for non-specialist users.

Last but not least, the critical step in all SPT experiments is the construction of a fusion between the protein of interest and the fluorescent tag. Occasionally, this results in an inactive protein. Therefore, the functionality of each fusion protein must be carefully verified. If the fusion is non-functional, the fluorescent tag can be placed at the other end of the protein or the length and nature of the linker can be adjusted to suppress the interference of the tag.

## Conclusion

SPT is a powerful method, allowing biochemical experiments to be performed in the native environment of living cells. When combined with perturbations such as protein mutations, deletions, or overexpression it can be used to gain deep mechanistic understanding of molecular pathways, and it has been applied not just to the field of DNA repair^2,3,8^, but also to study the stringent response^9^, transcription^1,10^ and translation^11^. SPT is becoming more and more popular, not only in the field of microbiology but also in the eukaryotic field^12^. Furthermore, the availability of user-friendly microscopes and analysis tools^13^ will pave the way for STP to become a standard tool in any laboratory.

## References

1 Stracy, M. et al. Live-cell superresolution microscopy reveals the organization of RNA polymerase in the bacterial nucleoid. Proceedings of the National Academy of Sciences of the United States of America, doi:10.1073/pnas.1507592112 (2015).

2 Uphoff, S., Reyes-Lamothe, R., Garza de Leon, F., Sherratt, D. J. & Kapanidis, A. N. Single-molecule DNA repair in live bacteria. Proceedings of the National Academy of Sciences of the United States of America 110, 8063–8068, doi:10.1073/pnas.1301804110 (2013).

3 Uphoff, S., Sherratt, D. J. & Kapanidis, A. N. Visualizing protein-DNA interactions in live bacterial cells using photoactivated single-molecule tracking. Journal of visualized experiments: JoVE, doi:10.3791/51177 (2014).

4 Betzig, E. et al. Imaging intracellular fluorescent proteins at nanometer resolution. Science 313, 1642–1645, doi:10.1126/science.1127344 (2006).

5 Manley, S. et al. High-density mapping of single-molecule trajectories with photoactivated localization microscopy. Nature methods 5, 155–157, doi:10.1038/nmeth.1176 (2008).

6 Stracy, M. et al. Single-molecule imaging of UvrA and UvrB recruitment to DNA lesions in living Escherichia coli. Nature communications 7, 12568, doi:10.1038/ncomms12568 (2016).

7 Persson, F., Linden, M., Unoson, C. & Elf, J. Extracting intracellular diffusive states and transition rates from single-molecule tracking data. Nature methods 10, 265–269, doi:10.1038/nmeth.2367 (2013).

8 Liao, Y., Schroeder, J. W., Gao, B., Simmons, L. A. & Biteen, J. S. Single-molecule motions and interactions in live cells reveal target search dynamics in mismatch repair. Proceedings of the National Academy of Sciences of the United States of America 112, E6898–6906, doi:10.1073/pnas.1507386112 (2015).

9 English, B. P. et al. Single-molecule investigations of the stringent response machinery in living bacterial cells. Proceedings of the National Academy of Sciences of the United States of America 108, E365–373, doi:10.1073/pnas.1102255108 (2011).

10 Bakshi, S., Siryaporn, A., Goulian, M. & Weisshaar, J. C. Superresolution imaging of ribosomes and RNA polymerase in live Escherichia coli cells. Molecular microbiology 85, 21–38, doi:10.1111/j.1365-2958.2012.08081.x (2012).

11 Sanamrad, A. et al. Single-particle tracking reveals that free ribosomal subunits are not excluded from the Escherichia coli nucleoid. Proceedings of the National Academy of Sciences of the United States of America 111, 11413–11418, doi:10.1073/pnas.1411558111 (2014).

12 Rhodes, J., Mazza, D., Nasmyth, K. & Uphoff, S. Scc2/Nipbl hops between chromosomal cohesin rings after loading. eLife 6, doi:10.7554/eLife.30000 (2017).

13 Sarkar, P., Switzer, A., Peters, C., Pogliano, J. & Wigneshweraraj, S. Phenotypic consequences of RNA polymerase dysregulation in Escherichia coli. Nucleic acids research 45, 11131–11143, doi:10.1093/nar/gkx733 (2017).

